# Bayesian and likelihood phylogenetic reconstructions of morphological traits are not discordant when taking uncertainty into consideration. A comment on Puttick *et al*

**DOI:** 10.1101/114793

**Authors:** Joseph W. Brown, Caroline Parins-Fukuchi, Gregory W. Stull, Oscar M. Vargas, Stephen A. Smith

## Abstract

Puttick *et al.* [1] performed a simulation study to compare accuracy among methods of inferring phylogeny from discrete morphological characters. They report that a Bayesian implementation of the Mk model [2] was most accurate (but with low resolution), while a maximum likelihood (ML) implementation of the same model was least accurate. They conclude by strongly advocating that Bayesian implementations of the Mk model should be the default method of analysis for such data. While we appreciate the authors’ attempt to investigate the accuracy of alternative methods of analysis, their conclusion is based on an inappropriate comparison of the ML point estimate, which does not consider confidence, with the Bayesian consensus, which incorporates estimation credibility into the summary tree. Using simulation, we demonstrate that ML and Bayesian estimates are concordant when confidence and credibility are comparably reflected in summary trees, a result expected from statistical theory. We therefore disagree with the conclusions of PEA and consider their prescription of any default method to be poorly founded. Instead, we recommend caution and thoughtful consideration of the model or method being applied to a morphological dataset.

## Comparing point estimates to consensus summaries

Puttick *et al.* (hereafter, PEA) [1] report that ML tree inference under the Mk model results in higher topological error than Bayesian implementations. However, this result is driven precisely by the comparison of maximum likelihood point estimates (MLE) to Bayesian majority-rule (BMR) consensus trees. MLE topologies are fully resolved, but this stems from the standard binary tree searching algorithms employed and not from an explicit statistical rejection of unresolved nodes. Therefore, individual MLEs may contain edges with negligible statistical support. On the other hand, consensus summaries, independent of phylogenetic method, may have reduced resolution as a product of uncertainty arising by summarization across conflicting sampled topologies. Thus, a direct comparison between a consensus tree (i.e., BMR) and a point estimate (i.e., MLE) is inappropriate. BMR topologies of PEA are more accurate simply because poorly supported conflicted edges were collapsed, while MLE topologies were fully resolved, even if poorly supported. While contrasting MLE and Bayesian maximum *a posteriori* (MAP) or maximum clade credibility (MCC) trees would be a more appropriate comparison of optimal point estimates, the incorporation of uncertainty is an integral part of all phylogenetic analysis. Therefore, comparison of consensus trees from Bayesian and ML analyses hold more practical utility for systematists. For these reasons, we argue that the results of PEA are an artefact of their comparison between fundamentally incomparable sets of trees.

## Confidence and credibility are fundamental to inference

To avoid drawing untenable conclusions, it is *de rigueur* of any statistical analysis to explicitly assess the robustness of an inference. Non-parametric bootstrap sampling [3] is the overwhelming standard in phylogenetic confidence estimation. PEA did not assess edge support in their ML estimates, stating that morphological data do not meet an underlying assumption of the bootstrap statistical procedure that “phylogenetic signal is distributed randomly among characters,” but provide no references to support the assertion. Non-parametric bootstrapping has been a staple of phylogenetic reconstruction for decades, including for the analysis of discrete morphological characters. Like Bayesian credibility estimation, bootstrapping estimates confidence by assuming that empirical data are a representative sample from an underlying distribution of characters evolving independently under a shared process [3]. As PEA note, the assumption of independence may often be violated. However, this violation is fundamentally problematic to model-based phylogenetics in general. Contrary to PEA, Bayesian and frequentist approaches to confidence estimation are similar in the sense that both provide distribution-based summaries of uncertainty, the sole distinguishing factor of Bayesian approaches is the incorporation of prior densities. Many of the concerns raised in relation to the bootstrap can thus also be shared by Bayesian approaches and should not preclude its use more generally. While there are concerns about the use and interpretation of the bootstrap [4], genetic datasets are routinely bootstrapped. Without additional information, it may be reasonable to assume that individual characters in a morphological matrix would be more independent than adjacent sites from the same gene (for which the interdependence among characters is far better understood). We thus dispute the assertion that bootstrapping is uniquely problematic for morphological data.

While Bayesian approaches estimate credibility intervals during parameter sampling,assessment is equally fundamental to likelihood analyses. In addition to the bootstrap, alternatives such as jackknifing and the SH-like test [5] are also implemented in popular software packages such as RAxML [6], one of the programs used by PEA. ML packages also frequently offer an option to collapse edges on a MLE tree that fall below some minimum threshold length. Use of any of these options would enable a more sensible comparison of likelihood and Bayesian reconstructions.

## ML and Bayesian comparisons incorporating uncertainty

To measure the effect of comparing BMR and MLE trees, we used the simulation code from PEA to generate 1000 character matrices, each of 100 characters on a fully pectinate tree of 32 taxa, as these settings generated the most discordant results in PEA. Each matrix was analyzed in both Bayesian and ML frameworks using the Mk+G model [2]. Bayesian reconstructions were performed using MrBayes v3.2.6 [7], using the same settings as PEA: 2 runs, each with 5 x 10^5^ generations, sampling every 50 generations, and discarding the first 25% of samples as burnin. As in PEA, we summarized each analysis with a BMR consensus tree (i.e. only edges with ≥ 0.5 posterior probability are represented). Likelihood analyses were performed in RAxML v8.2.9 [6]. For each simulated matrix we inferred both the MLE tree and 200 nonparametric bootstrap trees. Accuracy in topological reconstruction was assessed using the Robinson-Foulds (RF) distance [8], which counts the number of unshared bipartitions between trees. We measured the following distances from the true simulated tree: d_BMR_, the distance to the Bayesian majority-rule consensus; d_MLE_, the distance to the MLE tree; d_ML50_, the distance to the MLE tree which has had all edges with *<*50% bootstrap support collapsed. Finally, for each matrix we calculate D_MLE_ = d_MLE_ - d_BMR_, and D_ML50_ = d_ML50_ - d_BMR_. These paired distances measure the relative efficacy of ML and Bayesian reconstructions: values of D greater than 0 indicate that ML produces less accurate estimates (that is, with a greater RF distance from the true generating tree).

As demonstrated by PEA, MLE trees are indeed less accurate than BMR trees (Figure 1; D_MLE_), with MLE trees on average having an RF distance 17.6 units greater than the analogous Bayesian consensus distance. However, when collapsing MLE edges with less than 50% bootstrap support, Bayesian and ML differences are normally distributed around 0 (Figure 1; D_ML50_), indicating that when standardizing the degree of uncertainty in tree summaries there is no difference in topology reconstruction accuracy. These results support the argument that the original comparisons made in PEA of MLE and BMR trees are inappropriate. Depending on the level of uncertainty involved, an optimal point estimate from a distribution (e.g., MLE or MAP) may be arbitrarily distant from a summary of the same distribution. And so, the differences in MLE vs. BMR are not expected to be consistent.

**Figure 1:**
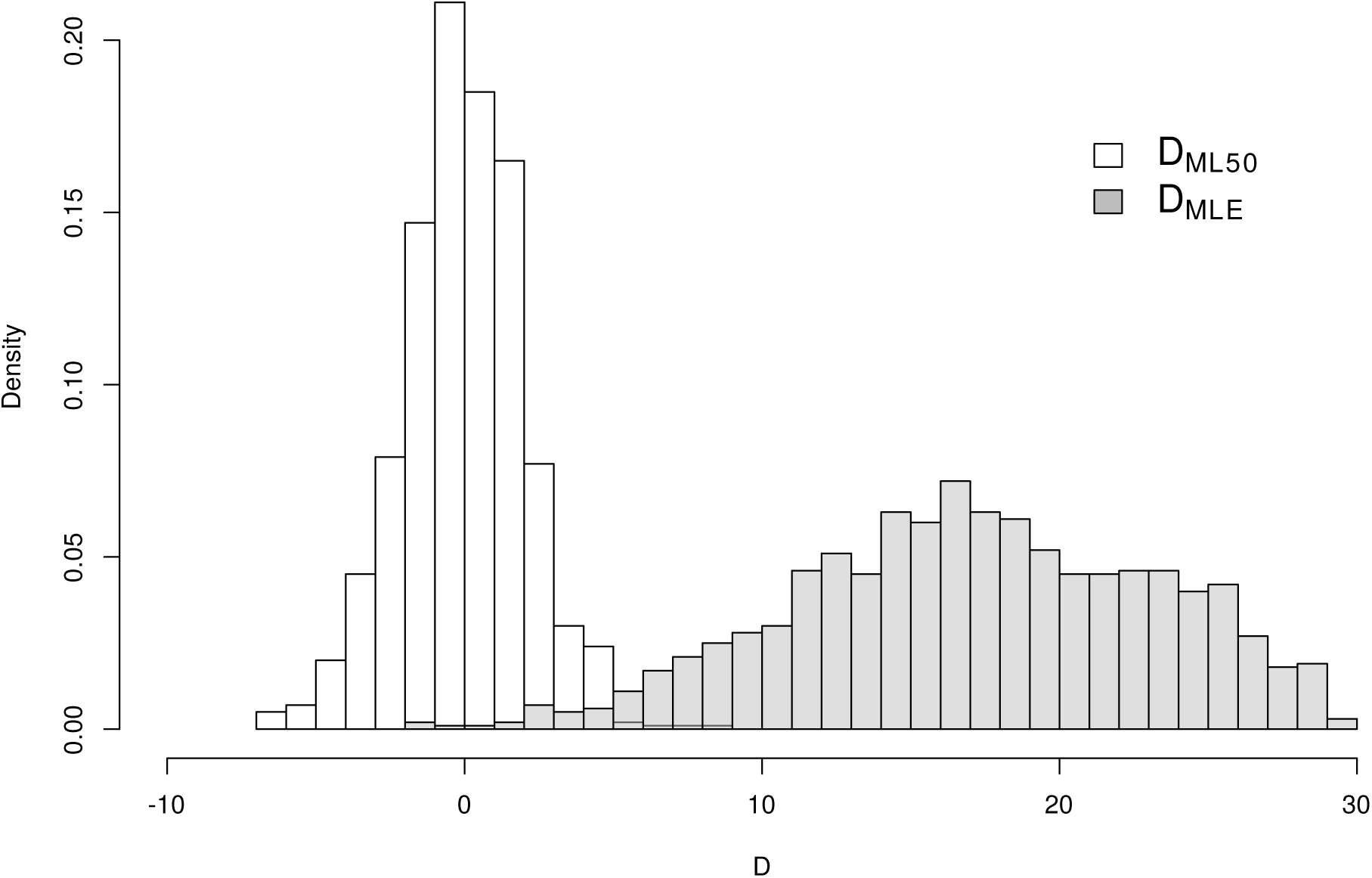
Topological accuracy of ML vs. Bayesian reconstructions for the most discordant comparison identified by PEA (see text). D measures how much larger ML distances are from the true tree (d_ML_) than are Bayesian distances (d_BMR_). MLE trees are indeed less accurate than BMRs (D_MLE_; mean = 17.63), but when conventional bootstrap thresholds are employed the difference in efficacy disappears (D_ML50_; mean = 0.43).

## The expected concordance of Bayesian and ML results

Our results reveal much greater congruence between Bayesian and ML estimates than suggested by PEA. This is to be expected and is reassuring. ML and Bayesian tree construction methods should yield similar results under the conditions in which they are often employed. While Bayesian tree reconstruction differs from ML by incorporating prior distributions, the methods share likelihood functions. In phylogenetics, researchers typically adopt non-informative priors, with a few exceptions (e.g., priors on divergence time parameters). Arguments can be made for pseudo-Bayesian approaches when care is taken to ensure that priors used are truly uninformative, which result in posterior probabilities that mirror the likelihood and are therefore congruent with ML [9, 10]. If prior distributions are formulated thoughtfully, as with [11] in shaping the Mk model using hyperpriors to accommodate character change heterogeneity, Bayesian methods can outperform ML. Alternatively, inappropriate priors can positively mislead [10]. Generally, when informative prior distributions are known or can be estimated using hierarchical approaches, Bayesian reconstruction methods may be strongly favoured over ML. It is unclear whether PEA intend to draw the comparisons discussed above as they do not describe any reasons to prefer Bayesian over ML in principle.

Although our results demonstrate general concordance between ML and Bayesian approaches when uncertainty is represented, further simulation work is needed to determine the extent and conditions of this concordance. Issues surrounding the application of Bayesian methods are particularly important in paleontology, where researchers often conduct inference upon very limited data. In these cases, it may be desirable to construct informative prior distributions when conducting Bayesian analyses [10]. The questions posed by PEA are sensible in light of current enthusiasm for statistical morphological phylogenetics. However, the relative performance of the implementations of the Mk model remain unresolved due to the authors’ misaligned treatment of confidence. This lack of resolution extends to their treatment of parsimony, which is invalid for the same reason as their ML comparison.

We do not advocate any one method for morphological phylogenetic reconstruction. Methods differ in model (Mk vs. parsimony), inferential paradigm (parsimony vs. ML/Bayesian),assumptions (prior distributions, model adequacy), interpretation, and means to incorporate uncertainty (ML/parsimony vs. Bayesian). We therefore recommend caution and thoughtful consideration of the biological question being addressed and then choosing the method that will best address that question. All inferential approaches possess strengths and weaknesses, and it is the task of researchers to determine the most appropriate given available data and the questions under investigation. The excitement of new morphological data sources and new means for analyzing these data should not overshadow the obligation to apply methods thoughtfully.

## Data accessibility

Simulated and inferred trees analyzed in this publication can be accessed in the electronic supplementary material.

## Authors’ contributions

All authors jointly developed the conceptual basis of the manuscript and study; J.W.B. conceived of and performed the simulations; J.W.B. and C.P.-F. drafted the manuscript; all authors contributed to the interpretation of results and the writing of the manuscript.

## Acknowledgements

We thank M. Puttick for an open and constructive discourse throughout this process. Two anonymous reviewers provided thoughtful reviews on an earlier draft of this manuscript. We thank Editor G. Carvalho and Associate Editor P. Makovicky for considering an appeal to an earlier decision on this manuscript. Finally, J.W.B. and C.P.-F. thank Annika Hansen for being a stalwart leading example of objective criticism. This is paper #1 of the PRUSSIA working group at UM.

